# Deep learning of the regulatory grammar of yeast 5’ untranslated regions from 500,000 random sequences

**DOI:** 10.1101/137547

**Authors:** Josh Cuperus, Benjamin Groves, Anna Kuchina, Alexander B. Rosenberg, Nebojsa Jojic, Stanley Fields, Georg Seelig

**Author notes:** equal contribution (Josh Cuperus, Benjamin Groves, Anna Kuchina, Alexander B. Rosenberg).

## Abstract

Our ability to predict protein expression from DNA sequence alone remains poor, reflecting our limited understanding of *cis*-regulatory grammar and hampering the design of engineered genes for synthetic biology applications. Here, we generate a model that predicts the translational efficiency of the 5’ untranslated region (UTR) of mRNAs in the yeast *Saccharomyces cerevisiae.* We constructed a library of half a million 50-nucleotide-long random 5’ UTRs and assayed their activity in a massively parallel growth selection experiment. The resulting data allow us to quantify the impact on translation of Kozak sequence composition, upstream open reading frames (uORFs) and secondary structure. We trained a convolutional neural network (CNN) on the random library and showed that it performs well at predicting the translational efficiency of both a held-out set of the random 5’ UTRs as well as native *S. cerevisiae* 5’ UTRs. The model additionally was used to computationally evolve highly translating 5’ UTRs. We confirmed experimentally that the great majority of the evolved sequences lead to higher translation rates than the starting sequences, demonstrating the predictive power of this model.

---

Precise control of protein expression is critical for cellular homeostasis and growth. One major layer of this control is exerted via post-transcriptional regulation of mRNAs, typically through the activity of their 5’ and 3’ untranslated regions. In *S. cerevisiae,* the effects of 5’ UTRs on translation have been characterized in detail for a few genes, pointing to the role of such features as upstream ORFs (Thireos et al. 1984; Werner et al. 1987), hairpins and other secondary structures (Linz et al. 1997; Yoon et al. 1992; Ringnér et al. 2005) and the Kozak sequence, i.e. the nucleotides (nt) immediately surrounding the AUG start codon (Hamilton et al. 1987). More recent studies have analyzed the functional consequences of polymorphisms and short sequence motifs (≤10 nt) in thousands or even tens of thousands of yeast (Dvir et al. 2013) and mammalian (Noderer et al. 2014) 5’ UTRs. However, this variation was targeted to nucleotides near the start codon, such that we are still unable to predict from sequence alone how the many distinct sequence and structural features of an entire 5’ UTR combine to regulate protein production.

A predictive model relating 5’ UTR sequence to protein production would not only provide novel insights into the grammar of biological *cis*-regulation, but it would also enable forward engineering of 5’ UTRs with tailor-made properties. Designing sequences with quantitatively predictable properties is a long-standing goal of synthetic biology and a prerequisite for accelerating the design-build-test cycle in metabolic engineering. Models have been designed, for example, to predict for *E. coli* the impact of a ribosome binding site on translation (Salis et al. 2009) or to understand how combinations of promoters and ribosome binding sites affect RNA and protein expression (Kosuri et al. 2013). However, so far, no generally applicable model has been generated that captures the effect of 5’ UTR sequence variation on protein production, primarily due to the lack of a dataset large and diverse enough to train such a model. Here, we overcome this limitation by using a library with 500,000 5’ UTR variants to generate a predictive model using a CNN.

While many different types of machine learning models have been applied successfully to biological data, CNNs in particular offer a combination of model power and interpretability, and as such have recently been used to predict transcription factor binding, DNase I hypersensitivity sites, enhancers, and sites of DNA methylation (Alipanahi et al. 2015; Zhou and Troyanskaya 2015; Kelley et al. 2016; Quang and Xie 2016; Lanchantin et al. 2016; Liu et al. 2016; Kleftogiannis et al. 2015; Wang et al. 2016). However, with yeast possessing only about 5,000 genes, measurement of the translational efficiency of this number of 5’ UTRs yields far too limited a dataset for accurate model building using CNNs. To generate data on a vastly larger scale, we designed a 5’ UTR library composed of completely random sequences of length 50.

With 4^50^ possible 5’ UTR sequences, the size of a resulting dataset of protein expression levels is limited only by experimental considerations and measurement capacity. The small number of nucleotides typically involved in the binding of proteins to RNA (4-8 nt) or in forming secondary structures (Weirauch et al. 2013) suggests that functional biological motifs will occur often and in a wide range of contexts within these random 5’ UTRs. Our study of alternative splicing corroborates the idea that highly predictive biological models can be learned from fully degenerate sequences (Rosenberg et al. 2015).

## Results

### 5’ UTR library and assay

Previous analyses of protein expression resulting from variants in a large library employed fluorescence activated cell sorting (FACS) for measurement (Dvir et al. 2013; Shalem et al. 2015; Lubliner et al. 2015; Noderer et al. 2014; Kosuri et al. 2013), wherein cells are separated into bins of differing fluorescence and the variants within each bin are sequenced. However, the FACS step limits the number of cells that can be assayed, thus also limiting the number of sequence variants that can be tested. To increase the number of 5’ UTRs that we could test simultaneously and to improve the resolution in measuring activity, we instead used a competitive growth assay based on the accumulation of the yeast His3 protein; the growth rate of cells in media lacking histidine is proportional to the level of their expressed His3 protein, a selection on continuous fitness values that is not reliant on arbitrary bins. In this selection, yeast are transformed with a library of plasmids carrying a *HIS3* reporter gene, each containing one of the random variants of the 5’ UTR sited immediately upstream of the start codon. The number of cells harboring each variant before and after growth in selection media is determined through sequencing, with the relative enrichment or depletion of a variant over time correlating with its *HIS3* expression. Since the number of cells in a selection is not limiting, a single culture can in principle be used to assay up to millions of variants. Similar growth-based selections have proven to be accurate in measuring activity differences (Starita et al. 2015; Rich et al. 2016).

We constructed a library of more than half a million 5' UTR variants (of which 489,348 were detected) (Fig. 1a, Online methods). With transcriptional regulation under the well-characterized low expression *CYC1* promoter and the *CYC1* terminator, (Chen et al. 1994; Guo et al. 1995; Martens et al. 2001; Watanabe et al. 2015; Yagil et al. 1998) and the use of a low copy number plasmid, the growth of each cell should reflect His3 protein accumulation. We performed a large-batch selection in media lacking histidine and supplemented with 1.5 mM 3-amino-1,2,4 triazole (3-AT, Supplemental Fig. 1a, see Methods), a competitive inhibitor of His3, collecting cells after ∼6.2 doublings. Using massively parallel sequencing, we quantified the relative change in abundance of each variant before and after selection.

**Figure 1.**
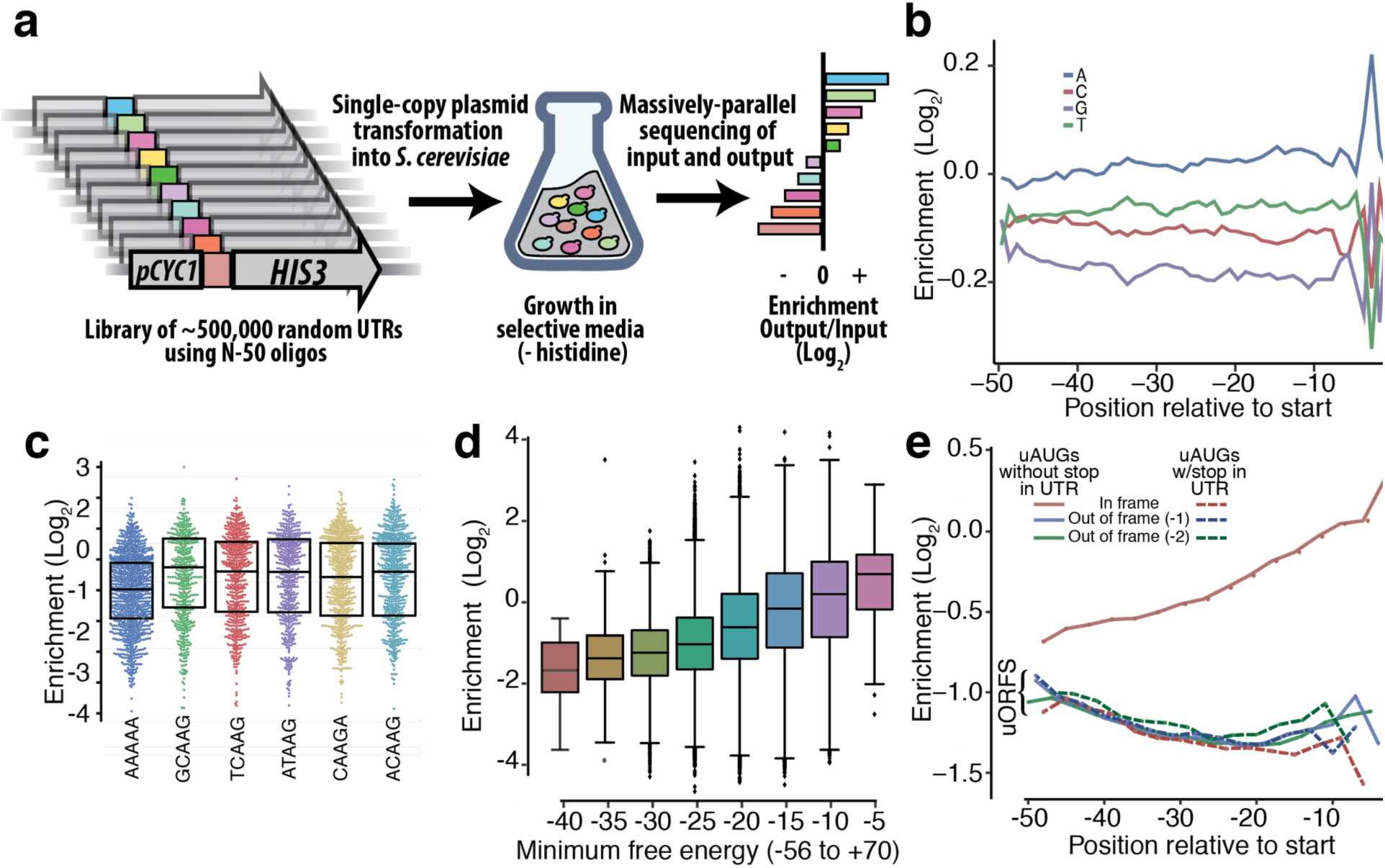
Experimental design and biological discovery. Experimental design of a liquid-based growth assay of 489,348 5’ UTR variants. Random 50 nucleotides were introduced directly upstream of the *HIS3* coding sequence, replacing the 56 nucleotides of the 5’ UTR of the *CYC1* promoter. These constructs were introduced into a low copy number plasmid, transformed into yeast without a native copy of *HIS3*, and competed in media lacking histidine. The enrichment of each UTR after growth was measured by using massively parallel sequencing before and after selection. (b) 5’ UTR enrichment scores per nucleotide were averaged at each position. (c) The Kozak sequences (-5 to −1 position) leading to the highest His3 protein expression compared to the most abundant yeast Kozak sequence (AAAAA). (d) The enrichment of 5’ UTRs based on the predicted minimum free energy of the −50 to +70 sequences. (e) The enrichment of 5’ UTRs based on the presence of an upstream AUG (uAUG) and a stop codon within the UTR. Upstream open reading frames (uORF) are characterized by an in-frame uAUG followed by a termination codon before the primary ORF start codon, or an out-of-frame uAUG followed by a stop codon before or after the primary ORF start codon.

To determine the accuracy of these pooled, competitive growth rate measurements, we chose 13 individual variants from the library with a range of translation levels and individually tested them. The relative growth rates of these 13 were similar to those measured in the bulk assay (R^2^= 0.84, Supplemental Fig. 1b, c). To further test the validity of our approach, we individually cloned 12 5’ UTRs from the library into a yellow fluorescent protein reporter, and measured fluorescence levels for these constructs using flow cytometry. We found good correlation between the data from the growth selection and flow cytometry (R^2^ = 0.61, Supplemental Fig. 1d), suggesting that results from the *HIS3* assay generalize to other gene contexts.

### Effects of 5’ UTR features

We next analyzed the effects of nucleotide identity at each position and in particular in the Kozak sequence, defined here as positions −5 to −1 relative to the start codon. Consistent with prior work (Dvir et al. 2013; Looman and Kuivenhoven 1993; Baim and Sherman 1988), the single nucleotide effects at positions −3 to −1 relative to the start codon were the most important, with an adenine in the −3 position the most beneficial to protein expression (Fig. 1b). This −3 preference for adenine is shared across many eukaryotes, including fungi, mammals and plants (Nakagawa et al. 2008). We examined the effect on protein expression of all possible Kozak sequences, as the library encompassed the 1,024 possible 5-mers at positions −5 to −1, with each 5-mer occurring on average in 478 different 5’ UTR contexts. Although the most common Kozak sequence for yeast is all adenine (Hamilton et al. 1987; Cavener and Ray 1991), we found that this sequence did not lead to the highest protein expression. In fact, 154 other Kozak sequences (122 of which contain an adenine at position −3) led to higher average protein expression than all adenine (the top 5 are plotted in Fig. 1c), contrary to the widely held belief that the most efficient Kozak is all adenine (supplemental Table 2). These highly efficient Kozak sequences are also present in the yeast genome in substantial numbers. Each of these top five Kozak sequences from our assay led to higher average translational efficiency than an all adenine sequence when assessed by ribosome profiling of native yeast genes (Supplemental Fig. 2a)(Pop et al. 2014).

We assessed the effect of secondary structure, which can influence ribosome initiation, scanning, and elongation (Pop et al. 2014; Rouskin et al. 2013). We first examined the correlation between the predicted minimum free energy (MFE) of the 5’ UTRs and protein expression. To calculate the predicted MFE, we used RNAfold (Gruber et al. 2015; Lorenz et al. 2011; Gruber et al. 2008) to fold each UTR sequence along with the 5’-most 70 nucleotides of the *HIS3* coding region. Binning the 5’ UTRs by their predicted MFE score, we found that lower MFE bins corresponded to decreased protein expression (Fig. 1d). Since the MFE provides only an aggregate measure of structure, we next looked at the effect of structure at each position in the 5’ UTR. We found that secondary structure had the largest effect on His3 expression when it occurs either near the 5’ end of the UTR or near the start codon (Supplemental Fig. 2b). 5’ secondary structure may reduce access to the 5’ cap by the 5’ cap binding complex, although only highly stable 5’ UTR secondary structures (<30 kcal/mol) markedly decrease eukaryotic translation rates (Babendure et al. 2006). Finally, as an alternative, simpler measure of secondary structure, we looked at the presence of hairpins with varying stem (5, 6, or 7 base pairs) and loop (0-25 nucleotides) lengths within the UTRs. We found that hairpins with longer stems and relatively short loops had the most negative impact on protein expression, perhaps because hairpins with longer loops form more slowly and are therefore scanned more readily by the translation machinery (Supplemental Fig. 2c). In spite of these correlations, secondary structure alone can explain only a small fraction of overall translational efficiency (MFE, enrichment correlation of R^2^ = 0.078, Supplemental Fig. 2d).

We analyzed the effects of upstream open reading frames (uORFs), characterized by an inframe uAUG followed by a termination codon before the primary ORF start codon, or an out-of-frame uAUG followed by a stop codon before or after the primary ORF start codon (Dvir et al. 2013; Wang and Rothnagel 2004; Morris and Geballe 2000). uORFs compete with the primary ORF, often producing nonsensical polypeptides and requiring translation to restart at the primary ORF start codon. Consistent with this competition, we found that the presence of a uORF led to greatly reduced protein expression (Fig. 1e, Supplemental Fig. 2e). On the other hand, a uAUG in-frame with the primary ORF, which results only in additional amino acids at the N-terminus of the translated protein, caused a minor reduction in expression. The effects of these in-frame uAUGs became more severe as the uAUG was located further towards the 5’ end of the UTR (Fig. 1e, Supplemental Fig. 2e), consistent with other reports (Rich et al. 2016; Dvir et al. 2013; Wang and Rothnagel 2004). The additional amino acids might cause a cumulative effect on translation, protein function or protein stability. Enrichment of 5’ UTRs with in-frame uAUGs correlated with the frequency with which the codons added upstream of the true AUG are used in *S. cerevisiae* (R^2^ = 0.75, Supplemental Fig. 2f), generally considered a measure of translational efficiency (Pop et al. 2014; Akashi 2003; Sharp and Cowe 1991).

### A convolutional neural network model

To better understand and engineer both native and synthetic UTR sequences, we sought to create a model capable of predicting our massive dataset. A comparison between different modeling approaches revealed several trade-offs. For example, a linear regression model with position-dependent 3-mer features (4^3^ x 48 = 3072 distinct features, R^2^=0.42) outperformed models with more complex but position-independent features (e.g. 6-mer model, 4^6^=4096 features, R^2^=0.33, Supplemental Fig. 3a). Given that many key features of translational efficiency in yeast have a position dependence—e.g. the identity of the nucleotide at position −3 or the frame of an upstream start codon—it is not surprising that a model that captures such position dependence can outperform a model that does not, even at the expense of using relatively simple features. However, we found that CNN models that capture not only position dependence but also non-linear interactions between features outperformed all simpler linear regression models on the task of predicting protein production from sequence alone.

CNNs typically consist of several layers of convolution that eventually feed into a classic feed-forward neural network. The first convolutional layer consists of many “filters” that essentially each learn a positional weight matrix (PWM). The output of this layer then feeds into further convolutional layers that can learn interactions between the different motifs recognized by each filter in the first layer. To choose the architecture of the model (such as the size of filters, number of filters and number of layers), we performed a hyper-parameter search using cross-validation on our training set. This search led us to choose a model with 3 layers of convolution, each with 128 filters of length 13. The convolutional layers then feed into a fully connected layer and finally a linear output layer. The output of our model is the predicted fitness score for each 5’ UTR, which should be proportional to translational efficiency.

In a test of the model with a held-out test set consisting of the top 5% of our library based on input read count, the CNN accounted for 62% of the observed variability (R^2^ = 0.62, Fig. 2a), outperforming a linear regression model trained on 3-mers as inputs (R^2^ = 0.42, Supplemental Fig. 3b). To understand the filters presented in the first layer, we scored 488,000 random 13-mers and created a PWM out of the top 1000 scoring sequences for each filter (Online methods). Twelve of the 128 filters in the first layer of the model learned uAUG motif variants, while eight learned motifs with stop codons (UAG/UGA/UAA) (Fig. 2b, Supplemental Fig. 4).

**Figure 2.**
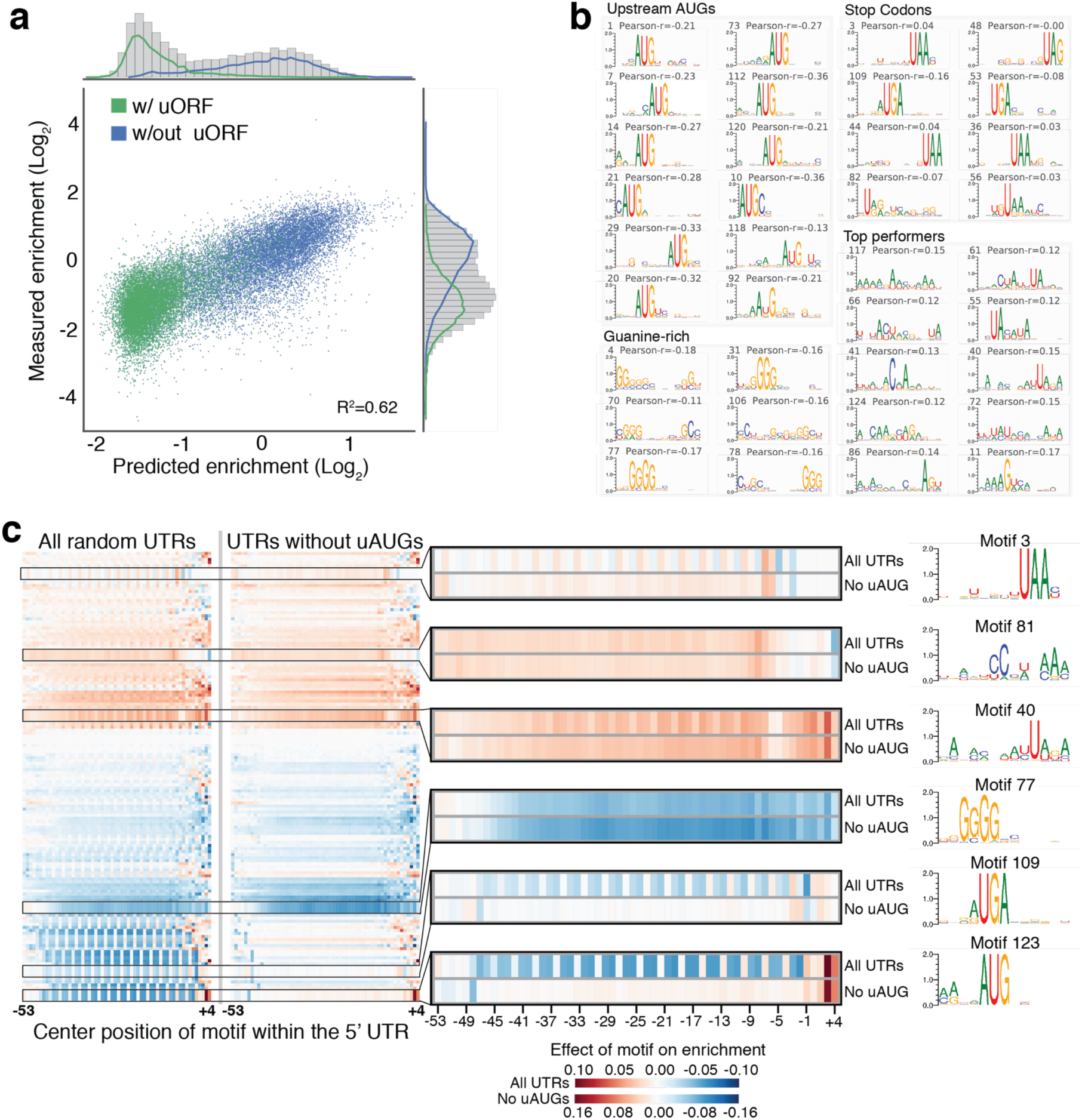
A convolutional neural network approach to model random 5’ UTR sequences. (a) A three-layer convolutional neural network model trained on random 5’ UTRs was tested on a heldout test set of the top 5% based on input read depth. Tested 5’ UTRs are specified by color for those with or without an upstream open reading frame (uORF). (b) 488,000 random 13-mers were scored for each filter in layer 1 of the CNN. The top 1000 13-mers were used to create a positional weight matrix (PWM) for each filter. These PWMs include motifs of start codons, stop codons, and guanine quadruplexes. Positive Pearson correlations indicate a positive effect on enrichment, while negative correlations indicate a negative effect on enrichment. (c)The effect of each motif per position was measured by assessing the Pearson correlation of motif score and enrichment at each position. Heatmaps of all 5’ UTRs (left), and those lacking upstream AUGs (right), including specific examples highlighting filters with different positional patterns are shown.

Additional filters resemble motifs involved in a G-quadruplex, an important motif in RNA secondary structure (Capra et al. 2010). However, several other motifs have no known function. They may correspond to the binding sites of RNA-binding proteins, as few binding sites for such proteins have been characterized in *S. cerevisiae* (Ray et al. 2013). Because the model should be interpreting not just translational efficiency, but features like RNA stability and changes in the transcriptional start site, filters could include several types of potential motifs (Fig. 2b, Supplemental Fig. 4).

Visualizing the positional dependencies of the first-layer motifs resulted in interpretable maps of the 5’ UTR sequence–function relationship. Some motifs had positional effects (Fig. 2c), such that they influenced protein expression differentially depending on their location within the 5’ UTR. Others showed a striking 3-nucleotide periodicity, reflecting their position relative to the reading frame of uAUGs. This periodicity was not present when 5’ UTRs lacking uAUGs were analyzed.

The second and third layer of the CNN can learn information about the interplay of lower-level filters. For example, some of the higher layers combine uAUG and stop codon filters to learn the concept of a uORF, as evidenced by the model predicting much lower protein expression for 5’ UTR sequences containing a uORF (see Fig. 2a). The model predicts that a 5’ UTR with only an in-frame upstream AUG will have a higher enrichment than one with an in-frame uAUG as well as an in-frame stop codon (Supplemental Fig. 3c). The model also predicts that a 5’ UTR with an in-frame uAUG as well as an out-of-frame stop codon will have only a small effect on expression (Supplemental Fig. 3c). Unlike a CNN, a simpler 3-mer model cannot capture these positional combinations.

Native 5’ UTRs may contain a higher density of motifs or higher order motifs not captured using a random library. We therefore asked whether the model could predict protein expression from native *S. cerevisiae* 5’ UTRs (Park et al. 2014). We constructed a library composed of 50 nt segments from all known native 5’ UTR sequences in the context of the *HIS3* reporter (Fig. 3a). Our model performed well on the task of predicting the impact of the native sequences (R^2^ = 0.60, Fig. 3b), giving us confidence that it captured the sequence information important for 5’ UTR function.

**Figure 3.**
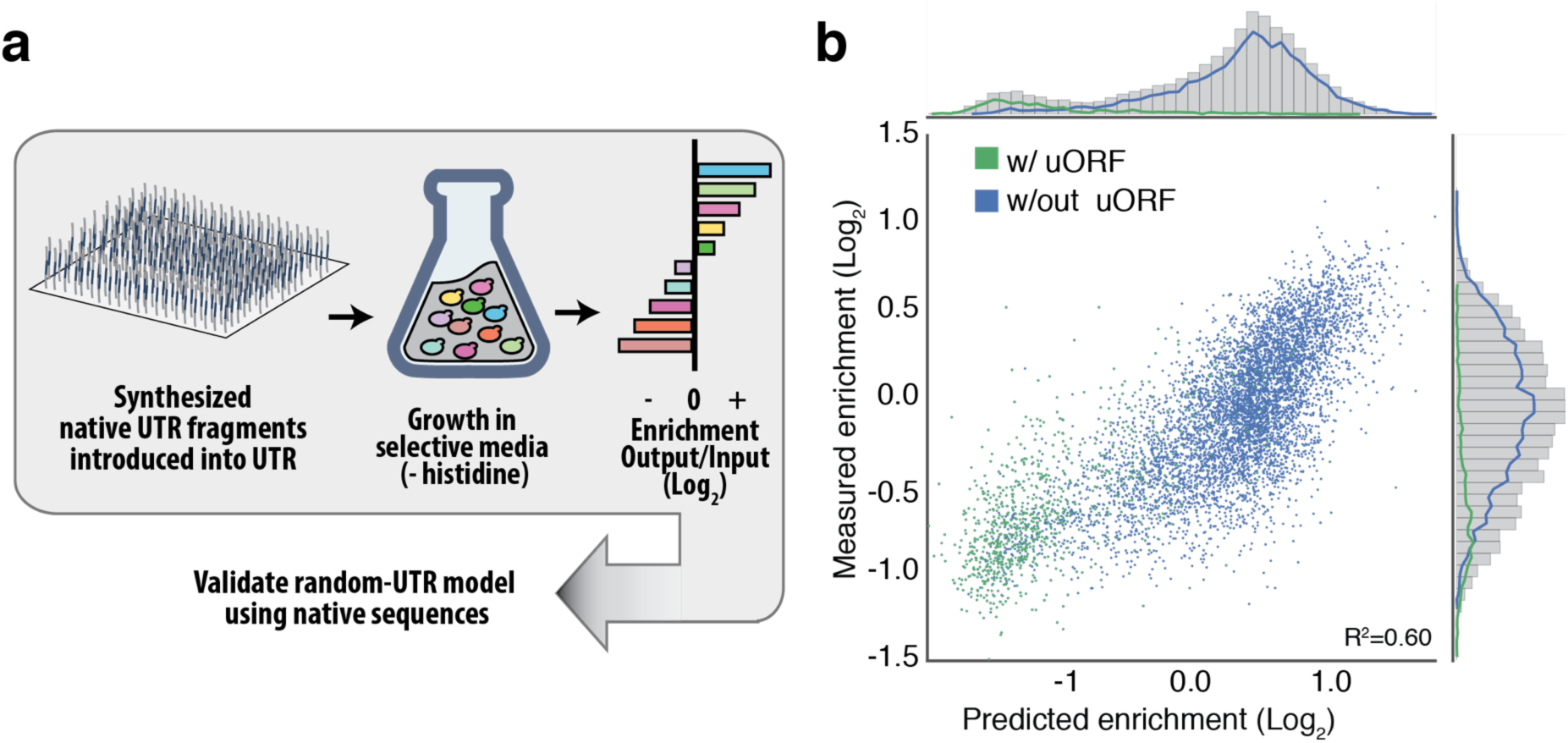
Validation of the CNN model on native 5’ UTRs. (a) Native 5’ UTR sequences were synthesized in 50 nucleotide fragments and introduced into the *HIS3*-based selection system. (b) Correlation of a native library with the predictions from our convolutional neural network built from random sequences.

### *In silico* evolution of 5’ UTRs

The design of functional sequences with user-defined properties is a compelling demonstration of the predictive power of a model. As a goal, we sought to use our model to improve the expression of a sample of random and native 5’ UTRs. We performed a model-guided *in silico* evolution of 200 5’ UTR sequences, half chosen from our random library and half from the native library, representing UTRs over the entire range of activity. During each step of the evolution, we made all possible mutations and selected the single nucleotide substitution predicted to result in the greatest increase in protein expression. We continued making changes until the predicted expression of each 5’ UTR plateaued (Fig. 4a and Supplemental Fig. 5a).

**Figure 4.**
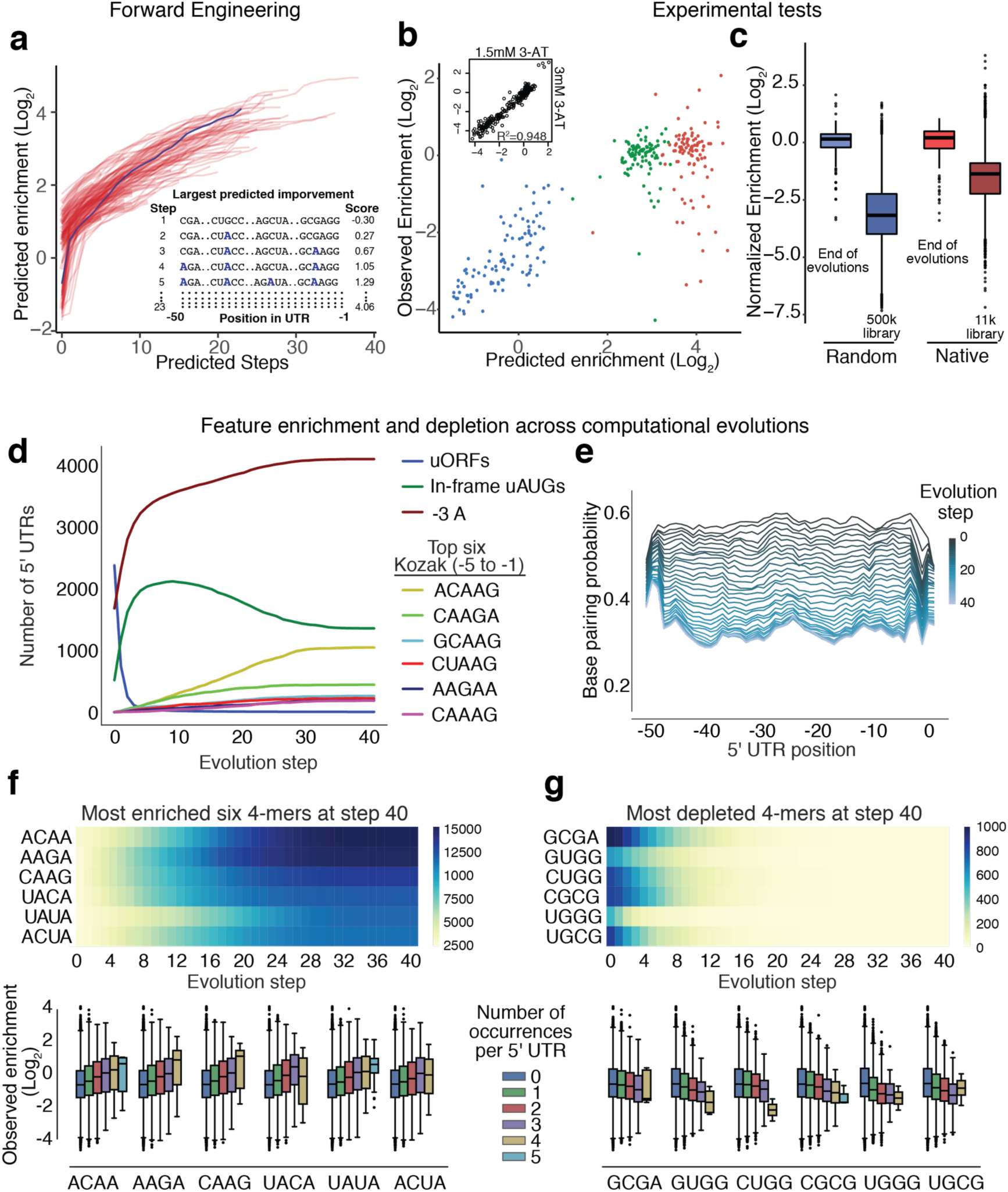
Model-guided optimization of 5000 random sequences. (a) Using our convolutional neural network, we iteratively predicted the optimal single nucleotide change in 100 random 5' UTR sequences until no additional increase in enrichment was predicted. An example of these changes can be seen in the inset. (b) The start, midpoint and endpoints from these evolutions were tested experimentally. The predicted and observed enrichments are plotted. (c) Experimental data from endpoints of the optimized 5' UTR sequences derived from both the random and native sets of sequence are compared to the enrichment distribution from the original random and native libraries. 5000 sequences from our random library were evolved over 40 steps and assayed for enrichment and depletion of common nucleotide features (d), secondary structure (e), and 4-mers (f and g). The effects of having the indicated enriched and depleted 4-mers occur multiple times in a single 5’ UTR were assessed by querying our original experimental random 5’ UTR dataset for enrichments based on the motif occurrences.

For 98 of the sequences derived from the random library and 93 from the native library, we were able to construct *HIS3* constructs for the starting, midpoint and endpoint of the evolutions. We then tested these 573 sequences in our growth assay. Our approach yielded improved expression for ∼94% and ∼84% of the sequences selected from the random and native libraries, respectively (Fig. 4b and Supplemental Fig. 5b). The relative expression due to these 5’ UTRs held in different 3-AT conditions (R^2^ > 0.93, Supplemental Fig. 6a). For the majority of sequences from both libraries, the largest increases in expression occurred between the starting and midpoint sequences, consistent with prediction from the evolutions. In both datasets, we also found that the degree to which the expression improved negatively correlated with the starting value, suggesting that it is easier to improve upon low expression 5’ UTRs than on high expression ones (random: R^2^ = 0.54; native: R^2^ = 0.86, Supplemental Fig. 6b). We also found that the majority of the endpoint sequences from the evolutions (88 out of 98 of the random library, and 75 out of 93 of the native library) performed better than 90% of their corresponding larger library after normalization (Fig. 4c, online methods).

To analyze where and how the CNN made changes, we expanded the number of random sequences that we computationally evolved to 5000, with each proceeding through 40 steps. We looked at the prevalence of simple characteristics, including uORFs, in-frame uAUGs, an A in the −3 position, favorable Kozak sequences and nucleotide bias (Fig. 4d, Supplemental Fig. 7a). The model selected against uORFs and structure (Fig. 4d, e and Supplemental Fig. 7b), and selected for an in-frame uAUG, A at the −3 position and overall A-rich composition except at positions −1 and −49, where Gs predominated (Supplemental Fig. 7a). Although one Kozak sequence (ACAAG) was the most prevalent, no single 5-nucleotide sequence dominated.

These more predictable changes were accompanied by more complex ones, revealed by analyzing the addition and removal of specific 4-mers (Fig. 4f, g and Supplemental Fig. 8). The 4-mers most enriched in the evolved sequences often appeared multiple times in a single 5’ UTR. We analyzed the experimental data collected from the random and native libraries and found additional copies of the enriched 4-mers correlated with continued increases in enrichment (Fig. 4f and Supplemental Fig. 9). Similarly, each additional copy of the depleted 4-mers correlated with reduced expression (Fig. 4g and Supplemental Fig. 9b). The spatial distribution of enrichment and depletion of all 4-mers was largely uniform across the 5’ UTRs, except at the ends of this region (Supplemental Fig. 8). The most enriched 4-mers and the most enriched Kozak sequence partially overlap with the reverse complement of part of the consensus motif for Nab3 (UCUUGU), a component of the transcription termination Nrd1 complex. The three 4-mers CAAG, ACAA and AAGA that match the reverse complement of the consensus site were highly enriched at the end of the evolutions compared to other motifs with the same nucleotide composition (p = 4.6 e-7, t-test, 42,906 occurrences in total for the three motifs), while 4-mers found within the motif itself (UCUU, CUUG and UUGU) occurred only 48 times at step 40.

## Discussion

Here we built and analyzed a library of approximately 500,000 random 5’ UTRs. We used the resulting data to train a CNN model that can predict the effect that any 5’ UTR in yeast will have on protein expression. Although the model was trained only on data from the random library, it performed equally well at predicting the behavior of native 5’ UTRs. This high quality of predictions is a direct result of the large training data set, compared to methods that consider only the limited set of approximately 5,000 native yeast 5’ UTRs.

Analysis and visualization of the model features allowed us to generate a comprehensive spatial map of the impact of *cis*-regulatory motifs on protein production. While some of the motifs identified were previously known and thus validated our approach, we also uncovered novel *cis*-regulatory motifs. For example, we identified 154 variants of the −5 to −1 region that outperformed the consensus Kozak sequence of five adenines. While functional roles for these motifs are supported by ribosome profiling data (Supplemental Fig. 2), the vast majority appear in the yeast genome in such low frequency that they could not be uncovered using native genes. Due to the large number of random sequences assayed in our library, we could observe these short elements in a large number of contexts, giving us power to identify these motifs and quantitate their impact.

We further demonstrated that our model can be used for the forward engineering of sequences with improved properties. Such computationally performed evolutions can dramatically reduce experimental overhead in the design of regulatory elements for synthetic pathways. Over the course of the model-guided sequence optimization, we found several motifs that became enriched. One of these, captured by the 4-mers CAAG and ACAA (Fig. 4d and f) and as the motif ACAAG in a convolutional filter (Fig. 2b), matches the reverse complement of the binding site for Nab3, a protein involved in transcriptional termination (Creamer et al. 2011; Porrua et al. 2012). We hypothesize that these sequences act to promote termination of transcription that occurs antisense to the *HIS3* reporter gene (Porrua and Libri 2015). Other motifs contain start or stop codons, G-quadruplex sequences known to influence expression or motifs of unknown function. Some of these motifs may represent target sites for RNA binding proteins, for which only a limited number of recognition sites have been identified to date in yeast 5’ UTRs.

Any approach that uses protein expression as its readout is potentially limited by its inability to distinguish among transcription, RNA processing and stability, and translation and protein stability. Transcription and post-transcriptional effects could be disentangled by direct measurement of RNA levels. Moreover, because our experimental approach relies upon growth selection, it is inherently less sensitive in detecting sequence variants that lead to poor protein expression. However, in any case, such variants are of limited interest at least for engineering applications.

Yeast has been the source of much of our knowledge of the highly conserved process of translation. Thus, we expect that our approach developed here will be similarly useful for understanding aspects of the biology of other organisms, for example allowing predictions about the impact of human genetic variation on transcription and translation (Dunham and Fowler 2013).

## Acknowledgments

This work was supported by DARPA under Contract No. W911NF-11-2-0068 to G.S., by the NSF through grant CCF 1317653 to G.S., and by NIH grant P41 GM103533 to S.F. S.F. is an investigator of the Howard Hughes Medical Institute.

## ONLINE METHODS

### Library construction

#### Synthetic 5’UTR library

We replaced a 56 bp *CYC1* 5’UTR fragment upstream of the *HIS3* ATG on a p415-CYC1 plasmid(Mumberg et al. 1995) with a library of 50 bp synthetic 5’ UTR fragments. The *CYC1* promoter is short (298 nucleotides), with well-established TATA-binding protein sites, upstream activating sequences (UASs) for *HAP1* (Pfeifer et al. 1987) and *MIG1* (Olesen et al. 1987; Treitel and Carlson 1995), and transcriptional start sites, and is regularly used as a consistent low expression promoter. The synthetic 5’ UTR fragments were constructed by annealing primers 126 and 127 containing an overlap region (ggacctttgcagca) and making the sequence double-stranded using the Klenow fragment of DNA polymerase I (NEB). The resulting fragment had a 50 bp random region and 60 bp and 33 bp 5’ and 3’ overlaps with the *CYC1* promoter and the *HIS3* coding sequence, respectively, including the ATG start codon. We inverse PCR-amplified the p415-CYC1 plasmid backbone with primers 132 and 133 using KAPA Hi-Fi polymerase (Kapa Biosystems), excluding the ATG start codon. Including the start codon in the library fragment served to prevent background plasmids not containing a library fragment from resulting in growth in media lacking histidine. The final library (YTLR200) was assembled using Gibson assembly^3^ and electroporated^4^ into 40 µl of 5-alpha electrocompetent *E. coli* (NEB) to yield 500,000 colonies.

#### Native 5’ UTR library

For the native library, we constructed 11,962 sequences representing native 5’ UTRs from the yeast genome^5^ in 50 bp fragments with 25 bp overlap if the UTR exceeded 50 bp in length, and in smaller fragments for UTRs shorter than 50 bp. 20 bp overhangs were added to both 5’ (acattaggacctttgcagca) and 3’ (ATGacagagcagaaagccct) ends of these sequences, again overlapping the *CYC1* promoter and *HIS3* gene on the p415-pCYC1 plasmid. The library sequences were purchased from CustomArray Inc. as a mixed oligo pool and amplified by qPCR using primers 126 and 142 in 15 cycles. The resulting fragment was assembled with the plasmid backbone via Gibson reaction and electroporated as described above, resulting in 200,000 colonies (YTLN200).

### Yeast transformation

For the library transformation into yeast, we followed the electroporation protocol described^6^. For the large synthetic 5’ UTR library (YTLR200), we used an overnight culture of BY4741(Baker Brachmann et al. 1998) diluted 1:50 into 50 ml of YPAD media(Amberg et al. 2005) and grown to OD 1.6. We prepared 400 µl of electroporation-competent cells as described and transformed with a mixture of 3.66 µg library plasmid YTLR200 linearized with EcoRI and 11.2 µg of DNA fragment PCR-amplified from YTLR200 with primers 134 and 135 to contain regions of overlap both upstream and downstream of the EcoRI restriction site. We grew the transformed library in 500 ml of synthetic dextrose media(Amberg et al. 2005) without leucine (SD-Leu) overnight and used colony counts from serial dilutions plated on SD-Leu to estimate library size. Using a longer region of homology (2.3 kb) led to improved transformation, resulting in ca. 2x10^6^ colonies. For the generation of the native 5’ UTR library (YTLN), the same protocol was followed. 6.7µg of EcoRI-digested library plasmid YTLN200 and 15.55 µg of PCR-amplified fragment (primers 134 and 135) were transformed into 800 µl of electrocompetent BY4741 yeast cells with similar efficiency as YTLR library described above. For the transformation of individual plasmids into yeast strains, we followed a lithium acetate method^7^.

### Growth rates measurements

Yeast cultures were grown overnight at 30 **°**C in 5 ml until saturated. In 96-well plates, cultures were diluted 1:20 in 200 µl volume of minimal selective media. The plates were shaken at 30° in media lacking histidine and leucine and with 3-amino-1,2,4 triazole (Brennan and Struhl 1980) (3-AT, Sigma) in a Synergy H1 hybrid reader (Biotek). Mean (n=6) maximum doubling rate was determined by measuring the maximum slope of O.D. 660 measurements over 6 points of measurement +/-standard error.

### Oligonucleotides and DNA sequencing

Oligonucleotides were obtained from Integrated DNA Technologies with standard desalting purification.

Sanger sequence and analysis was performed as described^8^. Deep sequencing of plasmid DNA was performed on an Illumina Nextseq after purifying plasmid DNA using the Zymoprep yeast plasmid prep II (Zymo Research) and PCR amplification for 12 to 20 cycles.

### Library selection

Cells from the input population were collected for sequencing and for back dilution into the selection medium (SD–His–Leu + 1.5 mM 3-AT) in triplicate adding 1 x 10^8^ cells to 1 L medium. Each replicate was cultured for 20 h to logarithmic phase (O.D. A660 = 1.0, 6 x 10^9^ cells), after which 3 x 10^8^ cells were collected for sequencing.

### Optimization of the dynamic range of the selection assay

To optimize the dynamic range of our selection assay, we compared the growth of two yeast strains, one harboring the *HIS3* construct with the native *CYC1* 5’ UTR and the other with a 5’ UTR containing a strong hairpin known to impair translation(Dvir et al. 2013; Lamping et al. 2013). In the presence of various concentrations of 3-AT, we found a maximal separation of growth rate between the two strains at 1.5 mM 3-AT (Supplemental Fig. 1a).

### Strains and media

Yeast experiments used the BY4741 strain*. pCYC1-HIS3* was cloned into the pRS series^10^ of yeast vectors with the *LEU2* nutrient marker (pRS415). To construct the plasmids harboring the individual synthetic and native 5’ UTRs, we designed a set of one forward and two reverse primers, each 30 base long with a 10 base overlap in the middle of the sequence for each sequence listed above. We added a 5’ acattaggacctttgcagca overhang to the forward primer (overlapping *CYC1* promoter), and either agggctttctgctctgtcat 3’ overhang (overlapping *HIS3* gene) or attcttcacctttagacat 3’ overhang (overlapping Venus gene) to the reverse primers. We obtained the oligos in a 96-well array (IDT), annealed them, filled them in with the Klenow fragment and cloned them into either the p415-pCYC1 backbone or p415-pCYC1-Venus backbone as described. The p415-pCYC1-Venus plasmid was constructed by replacing the *HIS3* sequence in the p415-pCYC1 plasmid used in our library construction with Venus via Gibson assembly.

### Enrichment analysis

For the random libraries, we first listed all identified UTRs. We then collapsed any sequences with a hamming distance of less than 3 and removed any with length less than 3. We used STAR (Dobin et al. 2013) to align reads from both our input libraries and selection libraries to this complete list of UTRs. Next, we counted the number of alignments to each UTR. To calculate the enrichment scores, we first added a “pseudocount” of 1 to the counts of each UTR in both inputs and selections and normalized the adjusted counts of each UTR by the total counts in each time point (input or selection), calculating the log enrichment of each sequence in the selection relative to the input. Native sequences were quantified similarly; however, because we started with known sequences, we were able to simply count the occurrences of each UTR in both the input and selection libraries as described above.

### Identifying features of 5’ UTRs

Using the enrichment scores derived from deep sequencing, we determined the average per-position score for each base, resulting in the plot in Fig. 2b. Ribsome profiling scores of native genes were calculated as the log-ratio of ribosome footprints counts over mRNA fragment counts. Minimum free energy was calculated using a window of −56 (the predicted transcriptional start site) and +70 using RNAfold (Gruber et al. 2015), then binned based on this MFE in increments of 5. Free base probabilities were also calculated using RNAfold. We searched for potential hairpins comprising combinations of hairpin length (5-7 nt) and loop length (0-24 nt), and then searched for perfect complementary pairs of 5-7 nt contained in a UTR. For each type of hairpin, we calculated the average enrichment scores of the subset of UTRs containing that type of hairpin. Plots were generated using matplotlib (Hunter 2007) or ggplot2 (Hadley Wickham 2009).

### Convolutional neural network training

All models were trained using the python package Keras (Chollet 2015). The test set was made from the 5’ UTRs with the most reads before selection (top 5%), using the rest of the data as a training set. All of our models were trained with the Adam optimizer (Kingma and Ba 2014) and early stopping was used to prevent overfitting to the training data. Cross-validation was performed on the training set to choose the model architecture. We tested combinations of the following hyper-parameters: convolutional filter width: [9, 13, 17, 25], number of convolutional filters per layer: [32, 64, 128, 256], number of convolutional layers: [2, 3, 4], number of dense layers: [1, 2], dropout probability in convolutional layers: [0, 0.15], dropout probability in dense layers: [0, 0.1, 0.25, 0.5], number of units in each dense layer: [32, 64,128, 256]. The best combination of hyper-parameters proved to be the following model architecture:

Layer 1: Convolutional, 128 filters (4x13), relu activation, 0.15 dropout probability

Layer 2: Convolutional, 128 filters (1x13), relu activation, 0.15 dropout probability

Layer 3: Convolutional, 128 filters (1x13), relu activation, 0.15 dropout probability

Layer 4: Fully connected layer, 64 hidden units, relu activation, no dropout

Layer 5: Linear output layer, 1 output unit

### K-mer models

The same training and test data were used to train linear regression models based on k-mer features. We trained models that simply used k-mer counts in each 5’ UTR as features as well as training models using k-mers at each position as features (*e.g.* for a 3-mer model there are 64 possible 3-mer sequences and 48 positions, leading to 3072 model weights; 3072=43 x 48). We cross-validated to choose optimal L2 regularization parameter for all k-mer models.

### Visualization and analysis of convolutional filters

To visualize each filter in the first layer of convolution, we scored 488,000 random 13-mers with each filter. We then used the top 1000 (0.2%) scoring 13-mers as input into the weblogo3 (Crooks et al. 2004) program to generate motifs. In order to calculate whether a given filter output associated with increases or decreases in protein expression we calculated the correlation across all 5’ UTRs between each filter output score at each position and the enrichment scores.

### Forward engineering of sequences

5,000 random sequences and 100 native sequences were analyzed for the single nucleotide change that led to the largest predicted change from our CNN model. This was done iteratively for 40 steps. From these, the start, midpoint and endpoint of 100 sequences from the random library and the 100 native sequences were chosen for synthesis. Endpoints were chosen based on the step at which no additional predicted enrichment was attained. Sequences were synthesized by oligonucleotide array (Custom array), introduced using Gibson assembly, and transformed into yeast. These yeast transformants were grown, collected and sequenced as before. Deep-sequencing data were analyzed using the Enrich package to assess enrichment of sequences (Rubin et al. 2016). To directly compare the evolved sequences with our larger random and native libraries, we determined the differences in enrichment scores of the starting point sequences (present in both libraries). We then normalized the rest of the larger libraries by the slope of these starting point scores to account for the differences **in** the strength of selection due to the differences **in** the sizes of the larger libraries compared to the evolutions.

## References

Akashi H. 2003. Translational selection and yeast proteome evolution. Genetics 164: 1291–303.

Alipanahi B, Delong A, Weirauch MT, Frey BJ. 2015. Predicting the sequence specificities of DNA-and RNA-binding proteins by deep learning. Nat Biotechnol, 33: 831–8.

Amberg DC, Burke D, Strathern JN, Burke D, Cold Spring Harbor Laboratory. 2005. Methods in yeast genetics: a Cold Spring Harbor Laboratory course manual. Cold Spring Harbor Laboratory Press.

Babendure JR, Babendure JL, Ding J-H, Tsien RY. 2006. Control of mammalian translation by mRNA structure near caps. RNA 12: 851–861.

Baim SB, Sherman F. 1988. mRNA structures influencing translation in the yeast Saccharomyces cerevisiae. Mol Cell Biol 8: 1591–601.

Baker Brachmann C, Davies A, Cost GJ, Caputo E, Li J, Hieter P, Boeke JD. 1998. Designer deletion strains derived fromSaccharomyces cerevisiae S288C: A useful set of strains and plasmids for PCR-mediated gene disruption and other applications. Yeast 14: 115–132.

Brennan MB, Struhl K. 1980. Mechanisms of increasing expression of a yeast gene in Escherichia coli. J Mol Biol 136: 333–338.

Capra JA, Paeschke K, Singh M, Zakian VA, Pringle T. 2010. G-Quadruplex DNA Sequences Are Evolutionarily Conserved and Associated with Distinct Genomic Features in Saccharomyces cerevisiae ed. G.D. Stormo. PLoS Comput Biol 6: 1000861.

Cavener DR, Ray SC. 1991. Eukaryotic start and stop translation sites. Nucleic Acids Res 19: 3185–92.

Chen J, Ding M, Pederson DS. 1994. Binding of TFIID to the CYC1 TATA boxes in yeast occurs independently of upstream activating sequences. Proc Natl Acad Sci U S A 91: 11909–13.

Chollet F. 2015. Keras.

Creamer TJ, Darby MM, Jamonnak N, Schaughency P, Hao H, Wheelan SJ, Corden JL. 2011. Transcriptome-Wide Binding Sites for Components of the Saccharomyces cerevisiae Non-Poly(A) Termination Pathway: Nrd1, Nab3, and Sen1 ed. N.J. Proudfoot. PLoS Genet 7: 1002329.

Crooks GE, Hon G, Chandonia J-M, Brenner SE. 2004. WebLogo: A Sequence Logo Generator. Genome Res 14: 1188–1190.

Dobin A, Davis CA, Schlesinger F, Drenkow J, Zaleski C, Jha S, Batut P, Chaisson M, Gingeras TR. 2013. STAR: ultrafast universal RNA-seq aligner. Bioinformatics 29: 15–21.

Dunham MJ, Fowler DM. 2013. Contemporary, yeast-based approaches to understanding human genetic variation. Curr Opin Genet Dev 23: 658–664.

Dvir S, Velten L, Sharon E, Zeevi D, Carey LB, Weinberger A, Segal E. 2013. Deciphering the rules by which 5 ′-UTR sequences affect protein expression in yeast. Proc Natl Acad Sci 110: E2792-801.

Gruber AR, Bernhart SH, Lorenz R. 2015. The ViennaRNA Web Services. In Methods in molecular biology (Clifton, N.J.), Vol. 1269 of, pp. 307–326.

Gruber AR, Lorenz R, Bernhart SH, Neubock R, Hofacker IL. 2008. The Vienna RNA Websuite. Nucleic Acids Res 36: W70–W74.

Guo Z, Russo P, Yun DF, Butler JS, Sherman F. 1995. Redundant 3’ end-forming signals for the yeast CYC1 mRNA. Proc Natl Acad Sci U S A 92: 4211–4.

Hadley Wickham. 2009. ggplot2: Elegant Graphics for Data Analysis. Springer-Verlag New York.

Hamilton R, Watanabe CK, De Boer HA. 1987. Compilation and comparison of the sequence context around the AUG startcodons in Saecharvmyces cerevisiae mRNAs. Nucleic Acids Res 15.

Hunter JD. 2007. Matplotlib: A 2D Graphics Environment. Comput Sci Eng 9: 90–95.

Kelley DR, Snoek J, Rinn JL. 2016. Basset: learning the regulatory code of the accessible genome with deep convolutional neural networks. Genome Res.

Kingma D, Ba J. 2014. Adam: A Method for Stochastic Optimization.

Kleftogiannis D, Kalnis P, Bajic VB. 2015. DEEP: a general computational framework for predicting enhancers. Nucleic Acids Res 43: 6.

Kosuri S, Goodman DB, Cambray G, Mutalik VK, Gao Y, Arkin AP, Endy D, Church GM. 2013. Composability of regulatory sequences controlling transcription and translation in Escherichia coli. Proc Natl Acad Sci U S A 110: 14024–9.

Lamping E, Niimi M, Cannon RD. 2013. Small, synthetic, GC-rich mRNA stem-loop modules 5’ proximal to the AUG start-codon predictably tune gene expression in yeast. Microb Cell Fact 12: 74.

Lanchantin J, Singh R, Lin Z, Qi Y. 2016. Deep Motif: Visualizing Genomic Sequence Classifications.

Linz B, Koloteva N, Vasilescu S, McCarthy JE. 1997. Disruption of ribosomal scanning on the 5’-untranslated region, and not restriction of translational initiation per se, modulates the stability of nonaberrant mRNAs in the yeast Saccharomyces cerevisiae. J Biol Chem 272: 9131–40.

Liu F, Li H, Ren C, Bo X, Shu W, Bulger M, Groudine M, Ong CT, Corces VG, Calo E, et al. 2016. PEDLA: predicting enhancers with a deep learning-based algorithmic framework. Sci Reports, Publ online 22 June 2016;| doi101038/srep28517 6: 250–257.

Looman AC, Kuivenhoven JA. 1993. Influence of the three nucleotides upstream of the initiation codon on expression of the Escherichia coli lacZ gene in Saccharomyces cerevisiae. Nucleic Acids Res 21: 4268–71.

Lorenz R, Bernhart SH, Höner zu Siederdissen C, Tafer H, Flamm C, Stadler PF, Hofacker IL. 2011. ViennaRNA Package 2.0. Algorithms Mol Biol 6: 26.

Lubliner S, Regev I, Maya L-P, Edelheit S, Weinberger A, Segal E. 2015. Core promoter sequence in yeast is a major determinant of expression level. Genome Res 25: 1008–1017.

Martens C, Krett B, Laybourn PJ. 2001. RNA polymerase II and TBP occupy the repressed CYC1 promoter. Mol Microbiol 40: 1009–19.

Morris DR, Geballe AP. 2000. Upstream open reading frames as regulators of mRNA translation. Mol Cell Biol 20: 8635–42.

Mumberg D, Müller R, Funk M. 1995. Yeast vectors for the controlled expression of heterologous proteins in different genetic backgrounds. Gene 156: 119–22.

Nakagawa S, Niimura Y, Gojobori T, Tanaka H, Miura K ichiro. 2008. Diversity of preferred nucleotide sequences around the translation initiation codon in eukaryote genomes. Nucleic Acids Res 36: 861–871.

Noderer WL, Flockhart RJ, Bhaduri A, Diaz de Arce AJ, Zhang J, Khavari P a, Wang CL. 2014. Quantitative analysis of mammalian translation initiation sites by FACS-seq. Mol Syst Biol 10: 748.

Olesen J, Hahn S, Guarente L. 1987. Yeast HAP2 and HAP3 activators both bind to the CYC1 upstream activation site, UAS2, in an interdependent manner. Cell 51: 953–61.

Park D, Morris AR, Battenhouse A, Iyer VR. 2014. Simultaneous mapping of transcript ends at single-nucleotide resolution and identification of widespread promoter-associated non-coding RNA governed by TATA elements. Nucleic Acids Res 42: 3736–3749.

Pfeifer K, Arcangioli B, Guarente L. 1987. Yeast HAP1 activator competes with the factor RC2 for binding to the upstream activation site UAS1 of the CYC1 gene. Cell 49: 9–18.

Pop C, Rouskin S, Ingolia NT, Han L, Phizicky EM, Weissman JS, Koller D. 2014. Causal signals between codon bias, mRNA structure, and the efficiency of translation and elongation. Mol Syst Biol 10: 770.

Porrua O, Hobor F, Boulay J, Kubicek K, D’Aubenton-Carafa Y, Gudipati RK, Stefl R, Libri D. 2012. In vivo SELEX reveals novel sequence and structural determinants of Nrd1-Nab3-Sen1-dependent transcription termination. EMBO J 31: 3935–48.

Porrua O, Libri D. 2015. Transcription termination and the control of the transcriptome: why, where and how to stop. Nat Rev Mol Cell Biol 16: 190–202.

Quang D, Xie X. 2016. DanQ: a hybrid convolutional and recurrent deep neural network for quantifying the function of DNA sequences. Nucleic Acids Res 44: e107–e107.

Ray D, Kazan H, Cook KB, Weirauch MT, Najafabadi HS, Li X, Gueroussov S, Albu M, Zheng H, Yang A, et al. 2013. A compendium of RNA-binding motifs for decoding gene regulation. Nature 499: 172–7.

Rich MS, Payen C, Rubin AF, Ong GT, Sanchez MR, Yachie N, Dunham MJ, Fields S. 2016a. Comprehensive Analysis of the SUL1 Promoter of Saccharomyces cerevisiae. Genetics 203: 191–202.

Rich MS, Payen C, Rubin AF, Ong GT, Sanchez MR, Yachie N, Dunham MJ, Fields S. 2016b. Comprehensive Analysis of the SUL1 Promoter of Saccharomyces cerevisiae. Genetics 203: 191–202.

Ringnér M, Krogh M, Mignone F, Gissi C, Liuni S, Pesole G, Hurowitz E, Brown P, Jansen R, Bashirullah A, et al. 2005. Folding Free Energies of 5′-UTRs Impact Post-Transcriptional Regulation on a Genomic Scale in Yeast. PLoS Comput Biol 1: e72.

Rosenberg AB, Patwardhan RP, Shendure J, Seelig G. 2015. Learning the Sequence Determinants of Alternative Splicing from Millions of Random Sequences. Cell 163: 698–711.

Rouskin S, Zubradt M, Washietl S, Kellis M, Weissman JS. 2013. Genome-wide probing of RNA structure reveals active unfolding of mRNA structures in vivo. Nature 505: 701–705.

Rubin AF, Lucas N, Bajjalieh SM, Papenfuss AT, Speed TP, Fowler DM. 2016. Enrich2: a statistical framework for analyzing deep mutational scanning data. bioRxiv.

Salis HM, Mirsky EA, Voigt CA. 2009. Automated design of synthetic ribosome binding sites to control protein expression. Nat Biotechnol 27: 946–950.

Shalem O, Sharon E, Lubliner S, Regev I, Maya L-P, Yakhini Z, Segal E. 2015. Systematic dissection of the sequence determinants of gene 3’ end mediated expression control. PLoS Genet 11: 1005147.

Sharp PM, Cowe E. 1991. Synonymous codon usage inSaccharomyces cerevisiae. Yeast 7: 657–678.

Starita LM, Young DL, Islam M, Kitzman JO, Gullingsrud J, Hause RJ, Fowler DM, Parvin JD, Shendure J, Fields S. 2015. Massively Parallel Functional Analysis of BRCA1 RING Domain Variants. Genetics 200: 413–22.

Thireos G, Penn MD, Greer H. 1984. 5’ untranslated sequences are required for the translational control of a yeast regulatory gene. Proc Natl Acad Sci U S A 81: 5096–100.

Treitel MA, Carlson M. 1995. Repression by SSN6-TUP1 is directed by MIG1, a repressor/activator protein. Proc Natl Acad Sci U S A 92: 3132–6.

Wang X-Q, Rothnagel JA. 2004. 5’-Untranslated regions with multiple upstream AUG codons can support low-level translation via leaky scanning and reinitiation. Nucleic Acids Res 32: 1382–1391.

Wang Y, Liu T, Xu D, Shi H, Zhang C, Mo Y-Y, Wang Z, Gardiner-Garden M, Frommer M, Cedar H, et al. 2016. Predicting DNA Methylation State of CpG Dinucleotide Using Genome Topological Features and Deep Networks. Sci Rep 6: 19598.

Watanabe K, Yabe M, Kasahara K, Kokubo T. 2015. A Random Screen Using a Novel Reporter Assay System Reveals a Set of Sequences That Are Preferred as the TATA or TATA-Like Elements in the CYC1 Promoter of Saccharomyces cerevisiae. PLoS One 10: e0129357.

Weirauch MT, Cote A, Norel R, Annala M, Zhao Y, Riley TR, Saez-Rodriguez J, Cokelaer T, Vedenko A, Talukder S, et al. 2013. Evaluation of methods for modeling transcription factor sequence specificity. Nat Biotechnol 31: 126–34.

Werner M, Feller A, Messenguy F, Piérard A. 1987. The leader peptide of yeast gene CPA1 is essential for the translational repression of its expression. Cell 49: 805–13.

Yagil G, Shimron F, Tal M. 1998. DNA unwinding in the CYC1 and DED1 yeast promoters. Gene 225: 153–62.

Yoon H, Miller SP, Pabich EK, Donahue TF. 1992. SSL1, a suppressor of a HIS4 5’-UTR stem-loop mutation, is essential for translation initiation and affects UV resistance in yeast. Genes Dev 6: 2463–77.

Zhou J, Troyanskaya OG. 2015. Predicting effects of noncoding variants with deep learning-based sequence model. Nat Methods.

